# Cytoplasmic male sterility and mitochondrial metabolism: evidence for low complex I contribution in male-sterile freshwater snail *Physa acuta*

**DOI:** 10.1101/2025.07.17.665361

**Authors:** Sophie Bererd, Damien Roussel, Sandrine Plénet, Loïc Teulier, Antoine Stier, Emilien Luquet

## Abstract

Cytoplasmic male sterility (CMS) is a well-known example of mitonuclear genetic conflict over sex determination in hermaphrodite plants, where mitochondrial genes maternally inherited sterilize the male function while biparental inherited nuclear genes restore it. CMS has been recently discovered in wild animals, in the freshwater snail *Physa acuta*. In this species, CMS is associated with two extremely divergent mitogenomes D and K compared to the classical mitogenome N. D individuals are male-steriles, while male fertility is restored in K individuals. We hypothesized that the extreme divergence of mitogenomes associated with CMS can impact mitochondrial aerobic metabolism, a necessary process in eukaryotic organisms by which energy is transduced from food to ATP. Our results suggest that CMS might be associated with an alteration of co-encoded complex I in D male-sterile individuals, although partly compensated by the nuclear-encoded complex II. K restored hermaphrodites have an unaffected complex I respiration, but exhibit higher mitochondrial proton leak and higher relative anaerobic contribution to cellular metabolism, suggesting an energy cost of bearing CMS and restorer genes, which could underlie the reduced growth of K mitotype. How complex I alteration might induce male-sterility remains to be determined, but could be linked to oxidative stress or a defect in ATP-synthesis rate.

## INTRODUCTION

The gene-centred view of evolution has revolutionised modern evolutionary biology, promoting that selection occurs at the level of the gene mediated by the phenotype (Williams, 1966). Conflict can arise when genes have opposing evolutionary interests, *i.e.* different strategies to maximize their own transmission, leading to antagonistic selective pressures over the phenotype (Werren, 2011). A fabulous example of genomic conflict impacting phenotype occurs in hermaphrodites, opposing nuclear and mitochondrial genes within individuals (Cosmides & Tooby, 1981; Werren & Beukeboom, 1998). Unlike nuclear genes transmitted by both parents, mitochondrial genes are usually inherited by one parent, most often the female, meaning that the fitness of mitochondrial genes only depends on the female function (Havird et al., 2019). This disparity in inheritance pattern between these genomes results in a conflict over sex determination (Chase, 2007; Cosmides & Tooby, 1981). Indeed, some selfish mitochondrial genes can emerge and sterilize the male function of hermaphrodites, who then become only functionally female. This phenomenon, known as cytoplasmic male sterility (CMS), increases the fitness of mitochondrial genes by enhancing their transmission though the female line (Budar et al., 2003; Chase, 2007; Saumitou-Laprade et al., 1994). In contrast, the male function sterilization leads to a loss of fitness for nuclear genes (Havird et al., 2019). Therefore, some nuclear genes neutralizing CMS (named restorer genes) can be selected to counteract sterilising effect of CMS genes by reverting individuals to hermaphroditism (known as restored hermaphrodite; Budar et al., 2003; Chase, 2007; Saumitou-Laprade et al., 1994). In natural populations, such a mitonuclear conflict results in a sexual polymorphism where male-sterile (female) individuals co-occur with hermaphrodite ones (normal and restored), a mating system called gynodioecy (Saumitou-Laprade et al., 1994). Gynodioecy, and the underlying dynamics of CMS and restorers genes in populations, is expected to evolve through a balance between selective advantage to be male-sterile (called female advantage, *i.e.* male-sterile individuals benefit from a better female fitness than hermaphrodites) and costs to bear CMS and restorers genes (Dufaÿ et al., 2007).

Gynodioecy was thought until recently to be a specificity of angiosperms (<1% of species, almost 20% families; Caruso et al., 2016; Dufay et al., 2014). Numerous studies highlight the association between divergent mitochondrial CMS lineages and the failure to produce pollen (review in Touzet & H. Meyer, 2014), suggesting that the functioning of mitochondria is at the core of the CMS mechanisms (*i.e.* male sterility, balance between female advantage and costs). Mitochondria are essential organelles as they are involved in various fundamental cellular functions, in particular cellular aerobic metabolism by transducing energy acquired from food to ATP (adenosine triphosphate) through oxidative phosphorylation (OXPHOS; Kadenbach, 2012; Logan, 2006; Vedel et al., 1999). The OXPHOS depends on the coordinated action of mitochondrial and nuclear genes because both encode for the proteins constituting the respiratory chain complexes and the ATP synthase (except the complex II that is fully encoded by the nuclear genome; Kadenbach, 2012). The divergence of CMS mitochondrial genomes may lead to altered functions through various and poorly known mechanisms (*e.g.* breakdown of coadaptation between mitochondrial and nuclear genomes, production of neoproteins by chimeric genes due to new open reading frames; Delph et al., 2007). In several CMS plant species, respiratory chain complex dysfunctions have been characterized and involve mainly complexes composed by protein subunits encoded by both mitochondrial and nuclear genes (Touzet & H. Meyer, 2014). For example, impairment of the complex I was described in *Nicotiana sylvestris* male-sterile mutants (Gutierres et al., 1997; Sabar et al., 2000) and in CMS transgenic rice (D. Tang et al., 2021). Cytochrome-c oxidase (COX - Complex IV) activity is also reduced by 50% in leaves of male-sterile wild beets (Ducos et al., 2001). Similarly, the activity of ATP synthase is reduced in CMS chilli pepper (Li et al., 2012). Understanding the physiological aspects of CMS is then crucial, as it helps to understand the mechanisms leading to male sterility and clarify the proximal causes underlying the evolutionary dynamics of gynodioecious systems.

Our study uses the freshwater snail *Physa acuta*, the recently-discovered and only known wild CMS animal model associated with naturally occurring mitochondrial types (David et al., 2022), allowing for the exploration of the mechanisms underlying CMS from the molecular to the fitness levels. Two mitochondrial genomes – named D and K mitotypes – have been associated with male sterility. These mitotypes are both extremely divergent from one another (mtDNA nucleotidic divergence ranged from 41% to 61%; Laugier et al., 2024) and from the classical mitochondrial genome, named N mitotype (35% to 57% between K and N and 28.6% to 53.2% between D and N; David et al., 2022; Laugier et al., 2024). D individuals are male-steriles (*i.e.* they produced a very low amount to no spermatozoa and showed a strong suppression of male reproductive behaviour; David et al., 2022) whereas K individuals are restored hermaphrodites (presence of nuclear restorer genes; Laugier et al., 2024). A recent study of Bererd et al. (2025) on the benefits and costs of CMS in *P. acuta* provided evidence that CMS is beneficial to female fitness in the absence of restorers (female advantage – D male-steriles have a better female fitness by producing higher number of eggs), while it is costly in their presence (K restored hermaphrodites have a lower growth rate). Interestingly, fitness benefits and costs were mediated by differences in body mass, suggesting that different growth and underlying metabolisms are involved. However, the whole-body metabolism was surprisingly similar among N, D and K mitotypes (Bererd et al., 2025). Altogether, these results suggested that the divergent mitochondrial genomes could cause impairments on cellular metabolism, altering the life history, fitness and fertility of organisms, but also that some compensation mechanisms may exist at the cellular level (*e.g.* higher number of mitochondria as proposed in CMS *Nicotiana sylvestris* by Sabar et al., 2000). The aim of the present study was thus to investigate how CMS and restorers genes were related to cellular metabolism potentially implicated in male sterility and balance between female advantage and costs in the *P. acuta* gynodioecious system (N normal hermaphrodite, D male-sterile, K restored hermaphrodite). We compared the male status (sterile or fertile), the growth rate, the mitochondrial aerobic metabolism and the anaerobic cellular metabolism between (i) individuals carrying CMS (D) and their non-CMS counterparts (N), and between (ii) individuals carrying CMS and K-specific restorers genes (K) and their non-CMS counterparts (N).

## MATERIAL AND METHODS

### Experimental design

*P. acuta* (Physidae, Hygrophila, Gastropoda) is a cosmopolitan widespread snail living in freshwater (Dillon et al., 2005). This species is a simultaneous hermaphrodite: sperm and oocytes are produced within the same organ called the ovotestis but sperm are stock in seminal vesicle for maturation, with a high preference for outcrossing (*i.e.* eggs are self-fertilised only when no mate is available to provide allosperm (Henry et al., 2005; Jarne et al., 2000; Tsitrone et al., 2003).

Three laboratory *P. acuta* lines were used: the mitotype N normal hermaphrodite, the mitotype D male-sterile and the mitotype K restored hermaphrodite. They were obtained from wild populations near Lyon (southeastern France) where the three mitotypes co-occurred and interbred, sharing a similar nuclear background (David et al., 2022; Laugier et al., 2024). In these populations, the prevalence of K-restorer genes in the nuclear background is high, leading to the restoration of male fertility in K snails (Laugier et al., 2024; Skarlou et al., 2025). Wild snails have been sampled and then reproduced in the laboratory (David et al., 2022; Laugier et al., 2024). The mitotype (N, D or K) of their offspring have been determined using a PCR test (see further details in Laugier et al., 2024). The three lines have been generated from these offspring and raised at 25°C (maturity in the laboratory is reached after 35-40 days at 25°C). N and K have been maintained as separate groups for 20 generations. D snails, as they were male-sterile, were fertilised by N snails through pair-crosses at each generation for 33 generations. The three lines share the same nuclear background, which carry restorer genes specific to K mitotype (effective in K lines but failing to restore CMS in D snails).

At 11 days post-hatching, 127 snails per mitotype were placed in a temperature-controlled room at 22.1 ± 0.1 °C and photoperiod 12h/12h. All the snails were reared individually in small boxes (90 ml) with dechlorinated tap water changed once a week and were fed with boiled and mixed lettuce twice a week.

### Male status and growth

At 34 days after the start of the experiment (45 days post-hatching), the male status (sterile or fertile) of each focal individual was assessed by pairing it to a virgin albino and by recording the pigmentation of the albino progeniture (unhatched offspring, about one week after the pairing using a binocular microscope): pigmented offspring indicated that the focal individual was male-fertile and on the contrary, no or albino offspring indicated male sterility (see David et al., 2022 for more details). The albino individuals are N mitotype, hermaphroditic snails (*i.e.* male-fertile and non-CMS), originating from the Montpellier Area (∼300km south of Lyon) and have been maintained as a large laboratory population for over 80 generations.

Before the pairing, whole-body mass (shell + body) was measured with a precision scale to the nearest 0.001mg. Maximal shell length was assessed using photography analysed with the software ImageJ (Schneider et al., 2012) at the nearest 0.001mm.

Note that only N and K male-fertile individuals (normal hermaphrodite and restored hermaphrodite, respectively) and D male-sterile individuals were selected for the comparisons of growth (whole-body mass and maximal shell length), mitochondrial aerobic metabolism and enzymatic activities.

### Mitochondrial aerobic metabolism

In order to compare the mitochondrial aerobic metabolism of the N, D and K mitotypes, 12-13 individuals per mitotype (total n = 37 individuals) were selected to assess mitochondrial functionality using high resolution respirometry (O2k, Oroboros; DatLab software; Makrecka-Kuka et al., 2015). Snails were anaesthetised on ice and dissected to sample the head and foot tissues (all the digestive and reproductive tissues were delicately removed). Tissue mass was measured with a precision scale at the nearest 0.001mg. Mechanical permeabilization of the tissue was performed by consecutive scissors cuts in 2 ml of a Mir05 solution (Salin et al., 2016) and verified as efficient in preliminary assays using additional chemical permeabilization with digitonin. Mitochondrial integrity was assessed during each run by the addition of 10 μM of cytochrome c, which had very little impact on respiration rate (+1.4% ± 1.3%), thereby proving the good integrity of our mitochondrial preparation. Four O2-k devices were used at an assay temperature of 21.5°C and a chamber volume of 2mL, with background calibration performed once before measurement sessions started, and 100% calibration performed daily. Mitotypes were randomly assigned to the four O2k devices, and were evenly distributed across five sessions of measurements. Mitochondrial aerobic metabolism was assessed using a relatively typical substrate-uncoupler-inhibitor titration (SUIT protocol; Pesta & Gnaiger, 2011; Fig.1).

**Figure 1.**
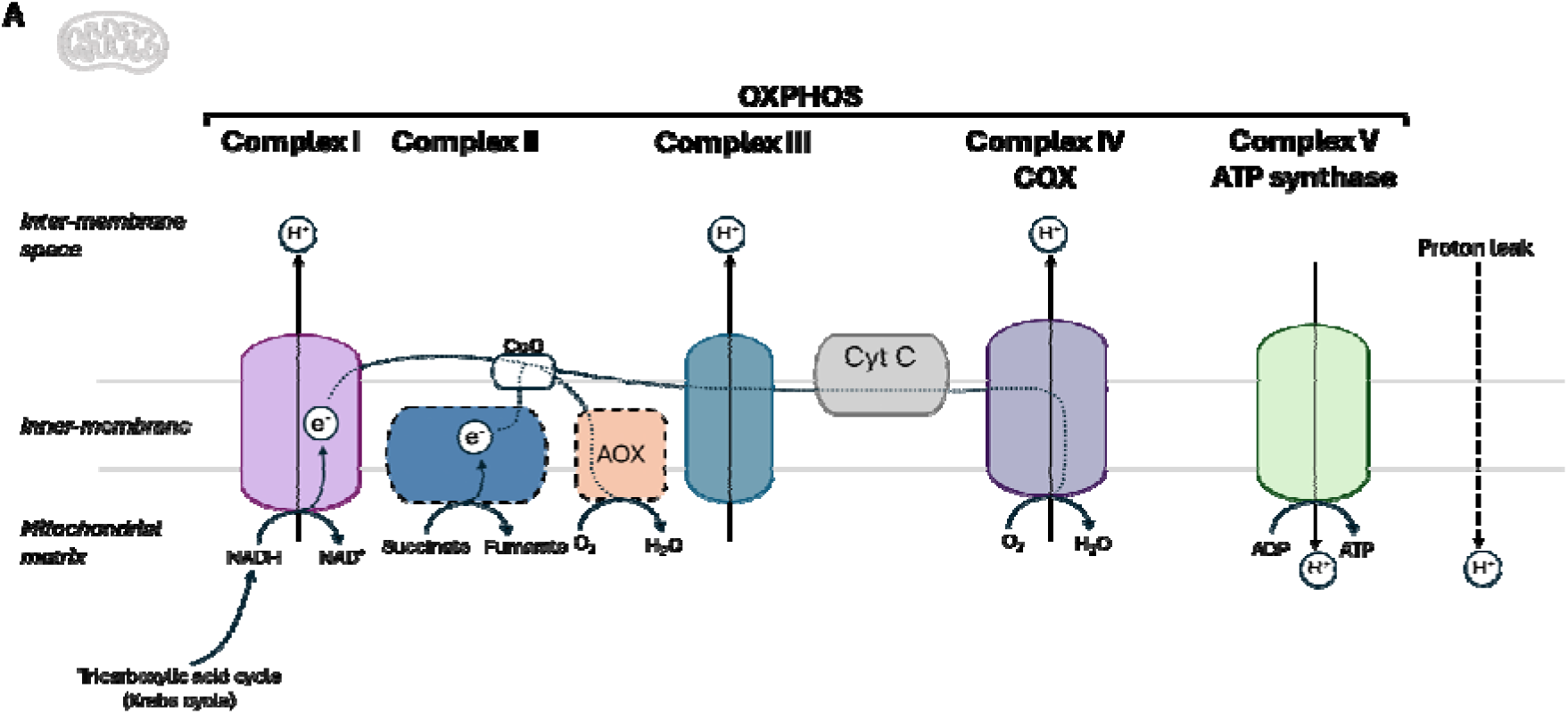
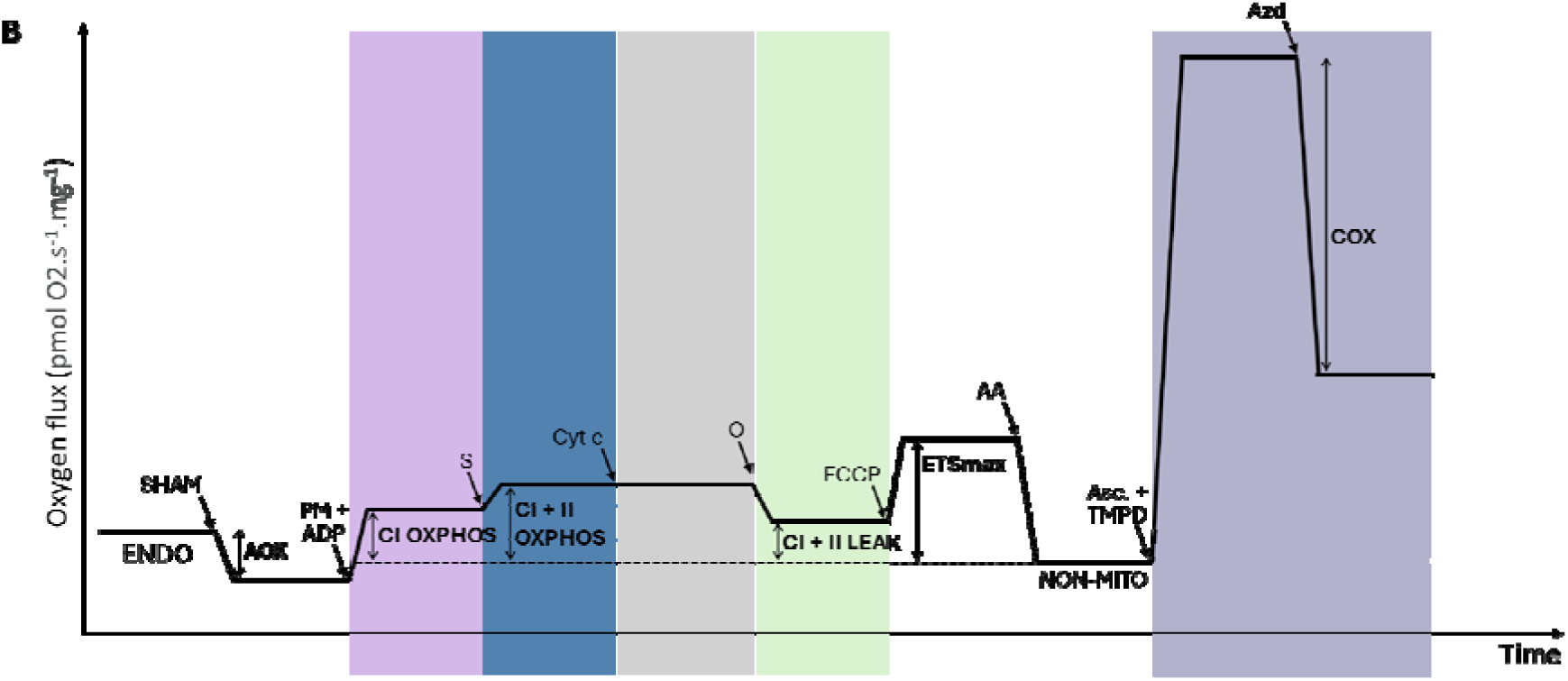
**A -** Schematic of the OXPHOS system. Mitochondrial and nuclear genes encode for the proteins constituting complexes I, III, IV and V (solid line border). Complex II and AOX are fully encoded by nuclear genes (black dashed border). The process relies on the respiratory chain formed by four protein complexes and the ATP synthase where a succession of redox reactions catalysed by four complexes (I, II, III and IV) creates a proton gradient, which is then used by complex V – ATP synthase – for the synthesis of ATP from ADP and inorganic phosphate. Proton Leak refers to the process where protons (H^+^) cross the inner membrane independently of ATP synthase. **B -** Graphical summary of the high-resolution respirometry substrate-uncoupler-inhibitor-titration (SUIT) protocol. *ENDO* respiration rate reflects the endogenous respiration of cells. *AOX* (alternative oxidase) respiration corresponds to the respiratory rate decrease in response to alternative oxidase inhibition by SHAM (Salicylhydroxamic acid or 2-Hydroxybenzohydroxamic acid). *CI OXPHOS* respiration rate fuelled by Complex I was measured after addition of its substrates Pyruvate and Malate with ADP (PM + ADP) and corresponds to the respiratory rate of the system with electron entry only through Complex I. *CI+II OXPHOS* respiration rate fuelled by Complexes I and II was measured after addition of Complex II’s substrates Succinate (S) and reflects the respiratory rate of the whole substrate-saturated system. *CI+II LEAK* respiration rate reflects the proton leak assessed by inhibiting Complex IV with Oligomicyn (O). *ETSmax* corresponds to the maximum capacity of the electron transport system and was measured after the titration of uncoupler FCCP (carbonyl cyanide p-trifluoromethoxyphenyl hydrazone). *COX* reflects the Complex IV maximum capacity to use O2. *NON-MITO* reflects the non-mitochondrial respiration rate, measured by the addition of antimycin A (AA).

During a pilot study aimed at setting up experimental conditions for high-resolution respirometry on *P. acuta* homogenate, we noticed surprisingly high levels of apparent non-mitochondrial respiration (*i.e.* after inhibition of complex III with 2.5, 5 or 10 μM of antimycin A). This residual respiration being sensitive to SHAM (Salicylhydroxamic acid or 2-Hydroxybenzohydroxamic acid), a known inhibitor of the alternative oxidase (AOX) pathway (Sluse & Jarmuszkiewicz, 1998), this suggests that AOX pathway is present and active in *P. acuta*. We thus systematically inhibited the AOX pathway at the beginning of the high-resolution respirometry assays with 1mM of SHAM, after stabilisation of endogenous cellular respiration. *AOX* respiration was thus calculated as the difference between endogenous and SHAM-inhibited respirations (Fig. 1B). Mitochondrial respiration was then fuelled by complex I substrates (5mM pyruvate and 2mM malate) and ADP was added at a concentration of 5mM to stimulate phosphorylating respiration (*CI OXPHOS*). Complex II substrate (succinate 10mM) was then added to achieve phosphorylating respiration driven by electron entry through both complexes I and II (*CI+II OXPHOS*). Cytochrome c 10 μM was added to verify mitochondrial integrity (see above). ATP synthesis was subsequently inhibited with 2.5 μM of oligomycin, leading to a non-phosphorylating respiration (*i.e.* driven by proton leak) referred to as *CI+II LEAK*. The uncoupler FCCP (carbonyl cyanide p-trifluoromethoxyphenyl hydrazone) was then titrated in 0.5 μM steps until maximum uncoupled respiration was achieved, which measures the maximum capacity of the electron transport system (*ETSmax*). Inhibition of complex III with antimycin A 2.5 μM led to mitochondrial respiration inhibition, and the residual (*i.e*. non-mitochondrial) O2 consumption left was subtracted from other respiration rates. Finally, electrons were fuelled directly to complex IV with 2mM ascorbate and 0.5mM TMPD (N,N,N’,N’-tetramethyl- p-phenylene-diamine), before inhibition of complex IV activity with sodium azide 100mM, the difference between these two respiration rates corresponding to complex IV maximum capacity to use O2 (*i.e.* COX respiration; Fig. 1B). All respiration rates were normalized per milligram of fresh tissue and expressed as pmolO_2_.s^-1^.mg^-1^ of tissue.

Mitochondrial respiration rates are sensitive to both mitochondrial quantity and intrinsic properties (*e.g.* functionality of electron transport). We also calculated three flux control ratios (*i.e.* ratios between different respiration rates) that are independent from mitochondrial quantity and thus purely reflect mitochondrial properties: 1. The efficiency of oxidative phosphorylation, *i.e. OXPHOS control efficiency*, calculated as: (*CI+II OXPHOS* - *CI+II LEAK*) / *CI+II OXPHOS (Gnaiger et al. 2020)*. This ratio gives the minimum proportion of O2 consumed contributing to oxidative phosphorylation; a ratio of 0.75 meaning that at least 75% of O2 consumption under phosphorylating conditions is dedicated to oxidative phosphorylation. *2.* The mitochondrial reserve to increase respiration from OXPHOS to maximal ETS respiration: *ETS reserve* calculated as (*CI+II ETS* - *CI+II OXPHOS*) / *CI+II ETS;* a ratio of 0.75 meaning that mitochondrial O2 consumption can increase by 75% in case of energy demand*. 3*. The relative contribution of complex I to maximal respiratory activity, when respiration is fuelled by both complex I and II substrates: *CI OXPHOS contribution*, calculated as 1 - ((*CI+II OXPHOS* - *CI OXPHOS*) / *CI+II OXPHOS);* a ratio of 0.75 meaning that complex I supports 75% of the mitochondrial O2 consumption when CI and CII substrates are provided together.

### Relative contribution of anaerobic metabolism to aerobic metabolism

The relative contribution of anaerobic metabolism to aerobic metabolism in the three mitotypes was measured by comparing enzymatic activities of the lactate dehydrogenase (LDH) and the citrate synthase (CS), respectively. LDH is a key enzyme of anaerobic metabolism, by converting pyruvate into lactate, it regenerates the NAD+ used by the glycolysis pathway to oxidize glucose-6-phosphate and synthesize ATP in the absence of oxygen (Emmett & Hochachka, 1981). CS is a key enzyme of mitochondrial aerobic metabolism as it is involved in the tricarboxylic acid cycle, which fuel the mitochondrial electron transport chain with reduced coenzymes (NADH and FADH2; Emmett & Hochachka, 1981; Krebs, 1970; Srere, 1971).

The head and the foot of 20 snails per mitotype (total n = 60 individuals) were sampled, weighted and frozen at −80°C. Tissues samples were thawed, then homogenised using a glass piston-glass Potter in 500μl buffer containing 100 mM sucrose, 50 mM KCl, 5 mM EDTA, 50 mM Tris-base, 1% fatty acid free BSA, pH 7.4. The resulting homogenates were kept at −80°C until measurements. After thawing, homogenates were vortexed and centrifuged at 1000 g for 10 min (4°C). The supernatants were retrieved and used for the LDH and CS enzyme assays. Enzyme activities were measured spectrophotometrically at 22°C. All results were expressed as nmol.min^-1^.mg^-1^ of tissue.

- **LDH activity** was measured in a reaction medium composed of 40mN imidazole (pH 7), 0.04% BSA and 200 µM NADH. Two replicates per homogenates were performed with 10μl and 20μl of supernatants and 250μl of the LDH reaction medium were laid in a microtiter plate. After 3 min incubation, the reaction was initiated by adding 4 mM pyruvate and the change in optical density at 340 nm was recorded for 3 min. The enzyme activity was quantified using an extinction coefficient of 6.22 mM^-1^.cm^-1^.
- **CS activity** was measured in a reaction medium composed of 100 mM Tris buffer (pH 8), 100 µM 5,5’-dithiobis(2-nitrobenzoic acid), 300 µM acetyl-CoA and 0.1% Triton-X-100. Homogenates were diluted ½ in a 100 mM phosphate buffer. Two replicates of 5μl and 7μl of diluted homogenates and 250μl of the CS reaction medium were deposited in a microtiter plate. After 3 min incubation, the reaction was initiated by adding 500 µM oxaloacetate and the change in optical density at 412 nm was recorded for 3 min. The enzyme activity was quantified using an extinction coefficient of 13.6 mM^-1^.cm^-1^.
- The **LDH/CS ratio** was calculated to estimate the contribution of anaerobic metabolism relative to aerobic metabolism and used as an index of relative glycolytic capacity. The technical repeatability (i.e. intra-class coefficient of correlation) of our enzymatic measurements based on duplicate was high, namely *R* = 0.94 for CS and *R* = 0.82 for LDH.

### Statistical analyses

#### Male status

A generalized linear model (GLM) following a binomial distribution was used to test the proportion of male sterility in the three mitotype by an analysis of variance (ANOVA).

#### Growth

The effect of the mitotype on whole body mass and on maximal shell length were tested by a linear model using an ANOVA.

#### Mitochondrial aerobic metabolism

A global linear mixed model was conducted to analyse **mitochondrial respiration rates** with “mitotype” and “state” as fixed factors (state corresponds to the specific mitochondrial respiration rates: AOX, CI OXPHOS, CI+II OXPHOS, CI+II LEAK, ETS, COX), the O2k chamber identity as a random factor to control for potential technical variations between the eight chambers used, “session (day of measurement)” as a random factor to control for technical variations between measurement sessions and a factor to account for identity of individuals. Linear mixed models (LMM) were conducted to analyse **mitochondrial flux control ratio** with “mitotype” as a fixed factor, the O2k chamber identity as a random factor to control for potential technical variations between the eight chambers used, and “session (day of measurement)” as a random factor to control for technical variations between measurement sessions.

The significance of fixed effects was tested using F-tests and the Kenward-Roger approximation for the degrees of freedom.

#### Enzymatic activities

Means of the two replicates for both LDH and CS activities were used in statistical analyses. LDH/CS ratio was calculated as the average of LDH means and CS means. Linear models and ANOVAs were conducted to analyse the effect of the mitotype on LDH activity, CS activity and LDH/CS ratio.

All statistical analyses were carried out in Rstudio 4.3.2 (R Core Team, 2023). Linear mixed models were performed using *lme4* package (Bates et al., 2015; p-values were obtained using *LmerTest* package, Kuznetsova et al., 2017). Post-hoc tests were performed using the *emmeans* package (Lenth, 2024). Tukey correction was used when the mitotype factor was significant (p < 0.05), and FDR adjustment was used specifically for LMM testing mitochondrial respiration rates considering the high number of pairwise comparisons.

## RESULTS

### Male status

Male sterility depended on mitotype (χ²_2, 337_ = 277.43, p < 0.001; Fig. 2). D individuals were almost all male-sterile as D had a much lower male-fertility proportion than N (normal hermaphrodite) and K (restored hermaphrodite; in logit scale, N – D: Est.±SE = 6.25 ± 0.79, z = 7.94, p < 0.0001; D - K: Est.±SE= −6.29 ± 0.79, z = 8.00, p < 0.001). N and K were mostly male-fertile and not significantly different from each other (in logit scale, N – K = −0.045 ± 0.467, z = −0.096, p = 0.995).

**Figure 2.**
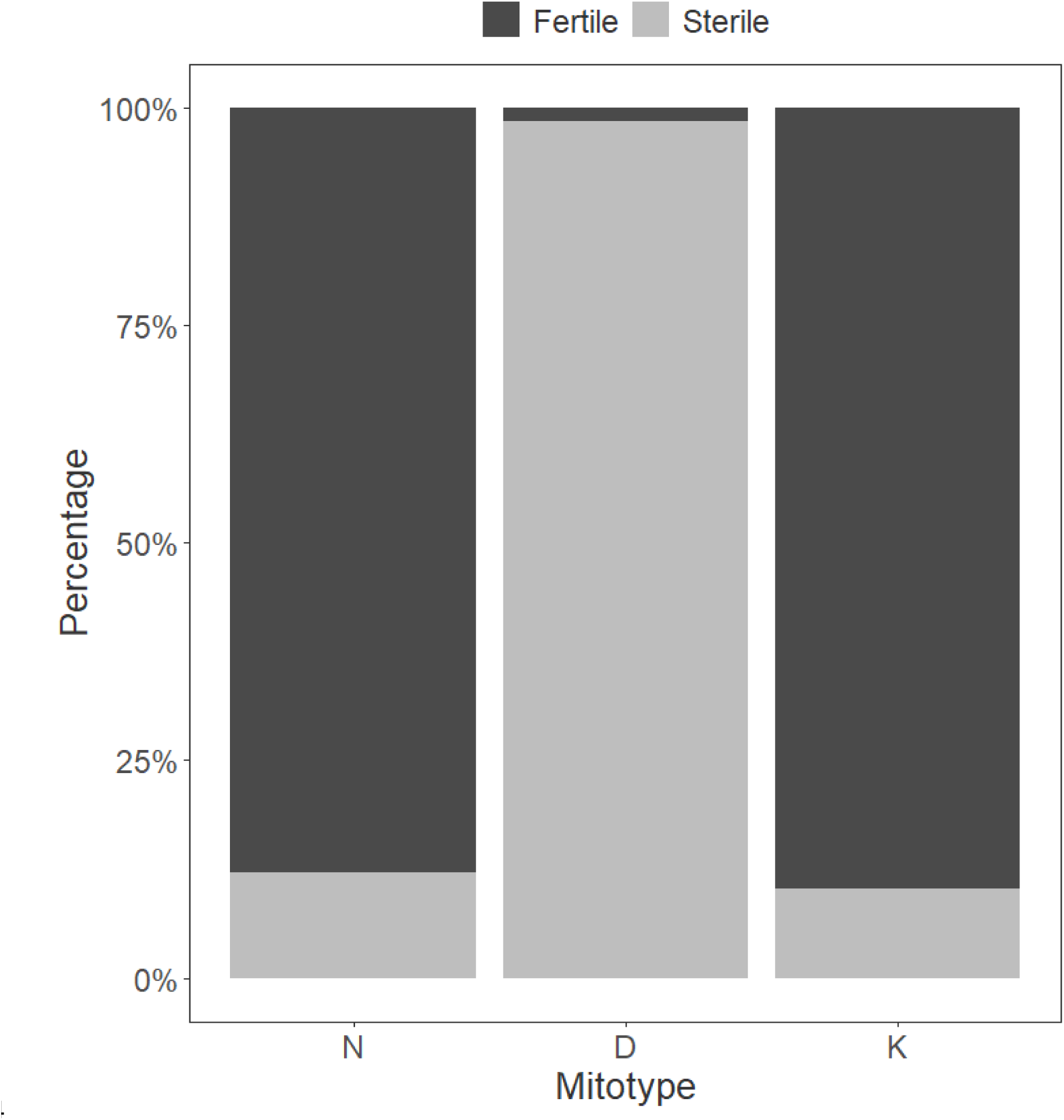
Percentage of male-fertile and male-sterile individuals in N, D and K *P.acuta* mitotypes. The total sample sizes were 124 N, 123 D and 127 K.

### Growth

**Whole-body mass** was significantly different between mitotypes (F_2, 341_ = 128.79, p < 0.001; Fig. 3A). D male-sterile snails were bigger than N and K male-fertiles (N - D: Est.±SE = −0.013 ± 0.003, t_341_ = −4.98, p < 0.001; D - K: Est.±SE = 0.042 ± 0.003, t_341_ = 15.78, p < 0.001). K had a significantly lower whole-body mass than N and D (N - K: Est.±SE = 0.029 ± 0.003, t_341_ = 10.46, p <0.001; Fig. 3A).

**Figure 3.**
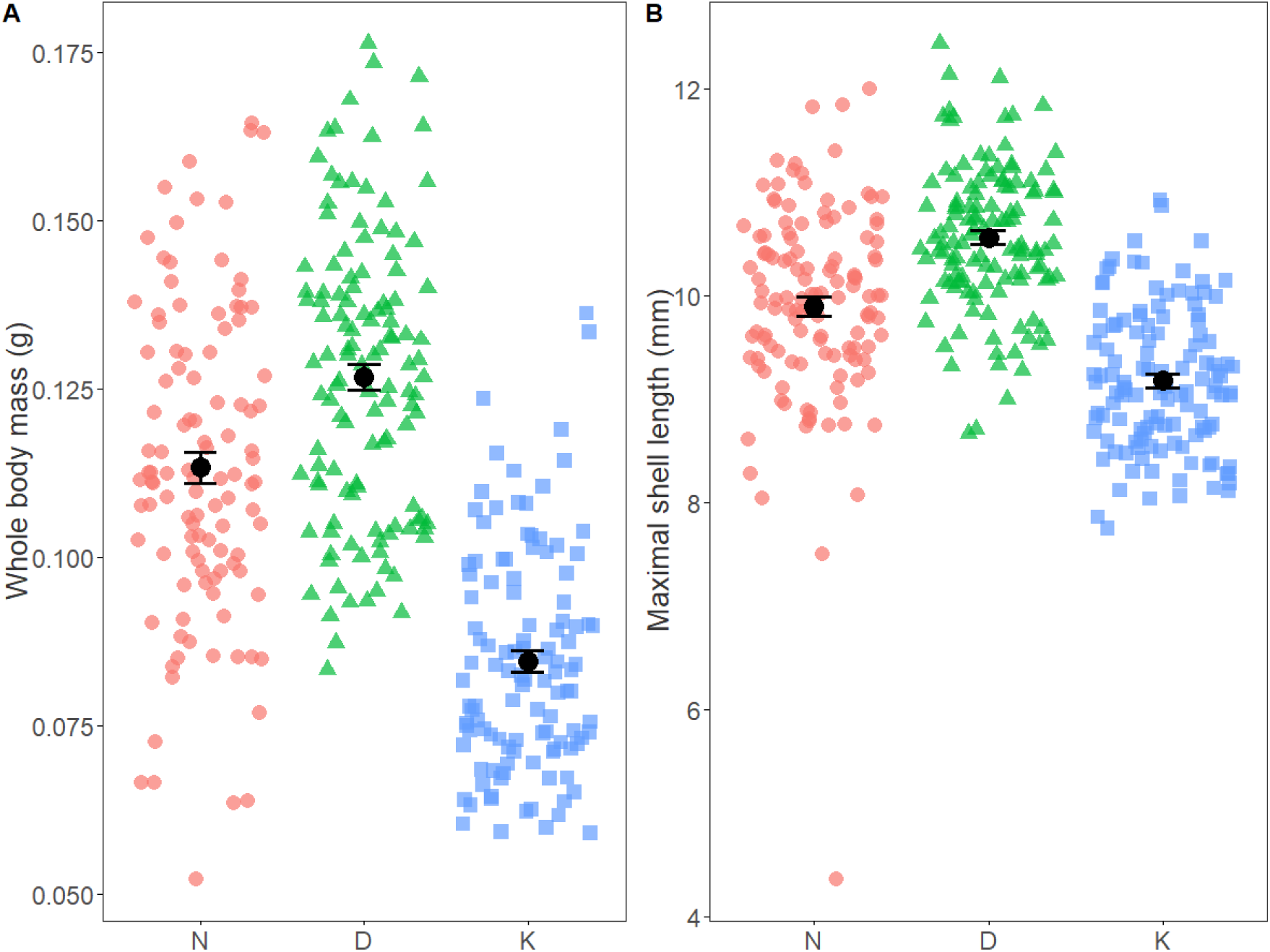
**A -** Whole-body mass (in g) for *P. acuta* normal hermaphrodites N (red circle), restored hermaphrodites K (blue square) and male-steriles D (green triangle). **B –** Maximal shell length (in mm). Black dots and black errors bars correspond to means ± standard error.

**Maximal shell length** was also significantly different between mitotypes (F_2, 338_ = 88.50, p < 0.001; Fig. 3B). D male-sterile had a longer shell length than N and K male-fertiles (N - D: Est.±SE = −0.662 ± 0.105, t_338_ = −6.30, p < 0.001; D - K: Est.±SE = 1.383 ± 0.104, t_338_ = 13.30, p <0.001). K had a significantly smaller shell length than N and D (N - K: Est.±SE = 0.720 ± 0.107, t_338_ = 6.75, p < 0.001; Fig. 3B).

### Mitochondrial aerobic metabolism Mitochondrial respiration rates

Mitochondrial respiration rates were significantly influenced by the mitotype*state interaction (interaction: *F*_10, 168.607_ = 2.33, p = 0.013; mitotype effect: *F*_2,9.779_ = 3.24, p = 0.083; respiratory state effect: *F*_5, 168.611_ = 255.56, p < 0.001). When looking for mitotype effect within each respiratory state, mitochondrial respiration rates related to oxidative phosphorylation through complex I and II combined (*CI+II OXPHOS*), maximal uncoupled respirations (*CI+II ETS*), and maximal oxidative capacity achieved under direct electron fueling to complex IV (*COX*) did not statistically differ between mitotypes (all p > 0.69; Fig. 4A, Tab. S1, Fig. S1). However, mitochondrial respiration through the alternative oxidase (AOX) pathway was lower in D mitotype compared to N and K (all p < 0.020, Tab. S1), a pattern also observed for mitochondrial respiration through complex I, although not fully significant (D - N: p = 0.079, D - K: p = 0.046). Finally, mitochondrial respiration rate related to proton leak through complex I and II (*CI+II LEAK*) was significantly higher in K mitotype than in N and D (Fig 4A; all p < 0.014, Tab. S1).

**Figure 4.**
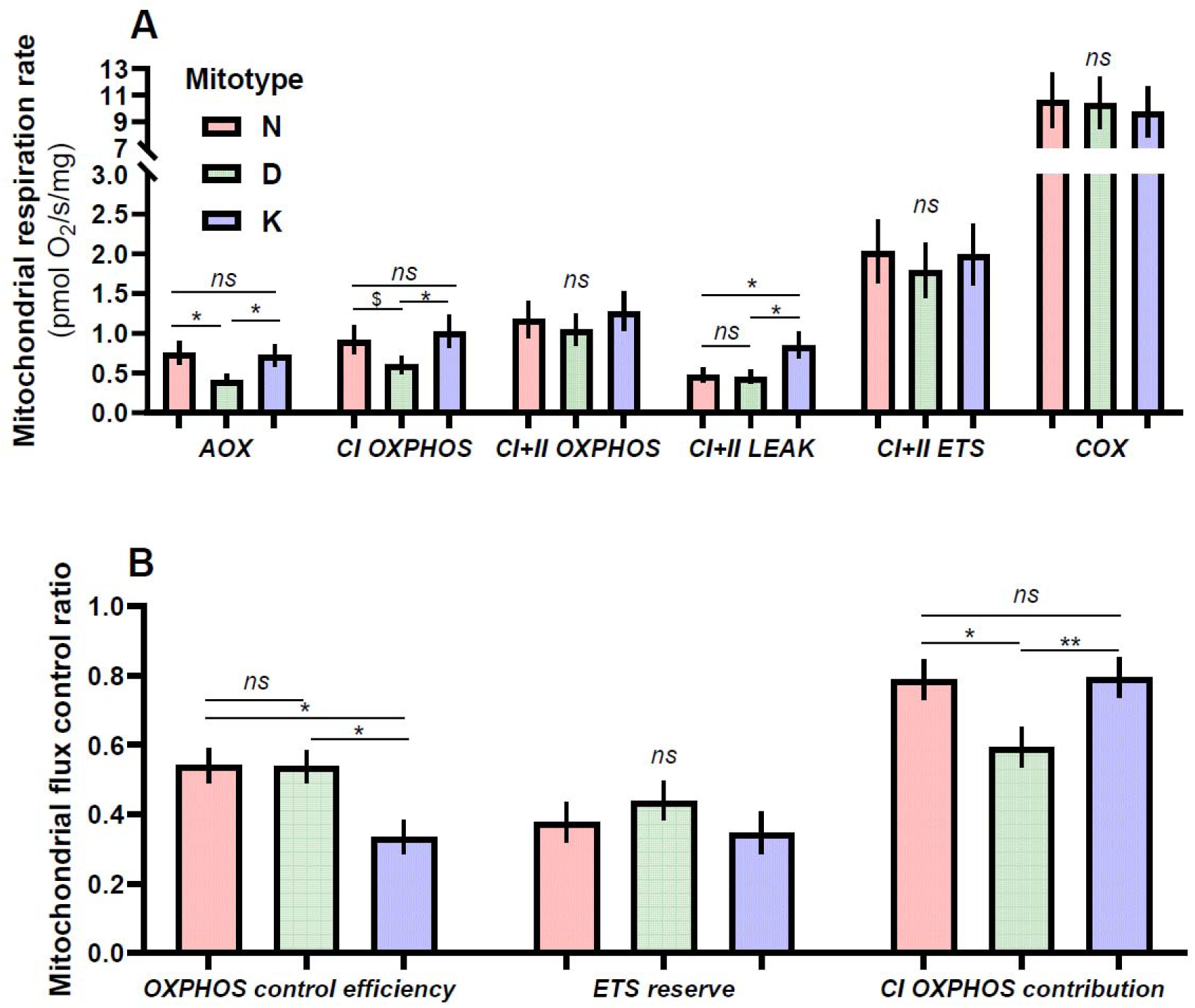
**A -** Mitochondrial respiration rates and **B –** Flux control ratios between mitotypes: N normal hermaphrodites (red), D male-steriles (green) and K restored hermaphrodites (blue) and. Figures are based on marginal means ± SE from statistical models and symbols indicate differences between mitotypes according to post-hoc tests (ns: p > 0.10, $: p < 0.10, *: p < 0.05, **: p < 0.01).

### Mitochondrial flux control ratios

The efficiency of the OXPHOS system (*OXPHOS control efficiency*) was significantly affected by mitotype (*F*_2,7.0744_ = 9.62, p = 0.009), with K mitotype having lower control efficiency than both N (*t*_6.01_ = 3.66, p = 0.025) and D mitotypes (*t*_7.19_ = − 3.48, p = 0.024; Fig. 4B). The ability to increase respiration from phosphorylating to uncoupled conditions did not significantly differ between mitotypes (*ETS reserve*: *F*_2,14.004_ = 0.78, p = 0.48). Finally, the contribution of complex I to OXPHOS respiration was markedly affected by mitotype (*F*_2, 29.977_ = 22.45, p < 0.001), with D individuals having lower contribution of complex I than N and K individuals (D - N: *t*_9.83_= −5.09, p = 0.013; D - K: *t*_5.91_ = −5.58, p = 0.004; Fig. 4B, Fig. S2).

### Relative contribution of anaerobic metabolism to aerobic metabolism

**LDH activity** was significantly influenced by the mitotype (F_2,57_ = 6.18, p = 0.004). D and K individuals had a significantly lower LDH activity than N (D - N: Est.±SE = 0.55 ± 0.17, t_57_ = 3.33, p = 0.004; K - N: Est.±SE = 0.44 ± 0.17, t_57_ = 2.65, p = 0.028). LDH activity in mitotypes K and D was not significantly different from each other (D – K: Est.±SE = −0.11 ± 0.17, t_57_ = −0.68, p = 0.78; Fig. 5, Fig. S3)

**Figure 5.**
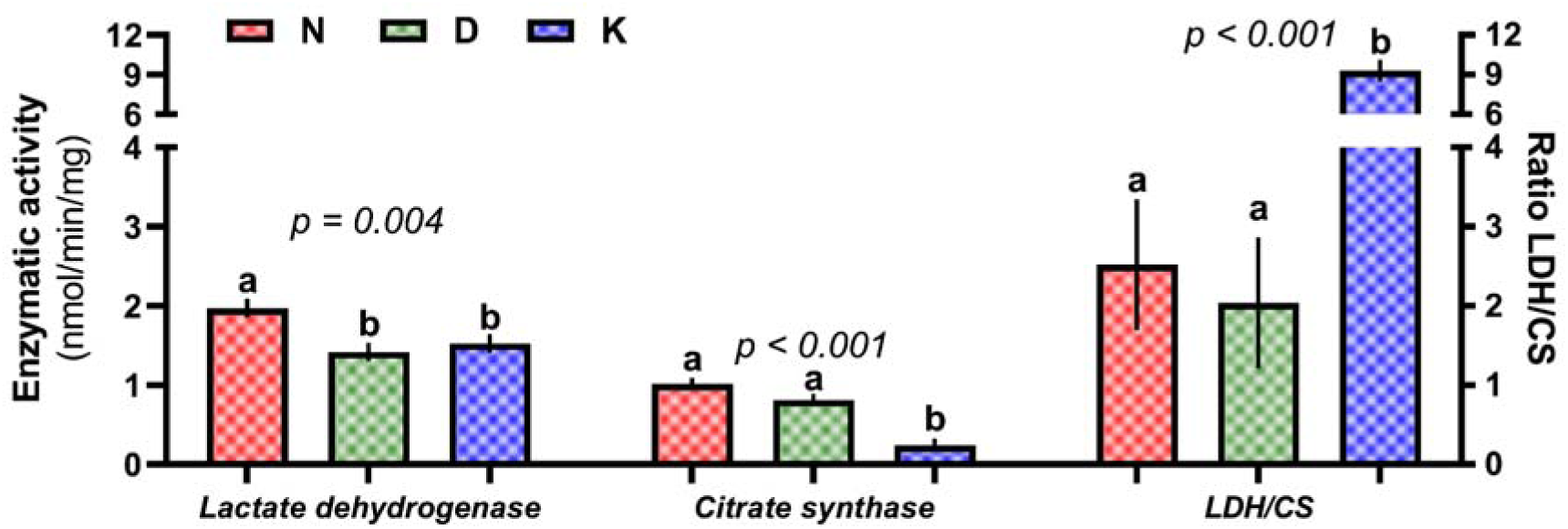
Enzymatic activities of lactate dehydrogenase (anaerobic metabolism) and citrate synthase (aerobic metabolism) of *P. acuta* N normal hermaphrodites (red), D male-steriles (blue) and K restored hermaphrodites (green). Means ± SE and different letters indicate significant differences according to ANOVAs and associated post-hoc tests.

**CS activity** was significantly influenced by the mitotype (F_2,57_ = 25.34, p < 0.001). K individuals had a significantly lower CS activity than N and D (N - K: Est.±SE = 0.77 ± 0.11, t_57_ = 6.86, p < 0.001; D - K: Est.±SE = 0.57 ± 0.11, t_57_ = 5.09, p < 0.001). CS activity in mitotypes N and D was not significantly different from each other (N – D: Est.±SE = 0.20 ± 0.11, t_57_ = 1.78, p = 0.19; Fig. 5, Fig. S3).

**The relative anaerobic contribution to metabolism (ratio of LDH activity to CS activity)** was significantly influenced by the mitotype (F_2, 57_ = 23.98, p < 0.001), being higher in K mitotype compared to N and D (N - D = Est.±SE = 0.47 ± 1.17, t_57_ = 0.41, p = 0.91; N - K: Est.±SE = −6.76 ±1.17, t_57_ = −5.78, p < 0.001; D - K: Est.±SE = −7.24 ± 1.17, t_57_ = −6.19, p < 0.001; Fig. 5, Fig. S3).

## DISCUSSION

Cytoplasmic male sterility (CMS) originates from a mitonuclear conflict where mitochondrial genes induce male sterility and nuclear genes restore male fertility in hermaphrodites. The dynamics of CMS and restorers genes is known to evolve through a balance between selective advantage to be male-sterile (*i.e.* female advantage) and costs to bear CMS and restorers genes (Dufaÿ et al., 2007). Male sterility and balance between female advantage and costs – key parameters of gynodioecy evolutionary dynamics – may be conditioned by changes in cellular metabolism. This study uniquely combined measurements of phenotype (growth trait as proxy of fitness advantage and costs) with an in-depth investigation of cellular metabolism mechanisms in the *P. acuta* gynodioecious system (N normal hermaphrodite, D male-sterile, K restored hermaphrodite; David et al., 2022; Laugier et al., 2024). Our results suggest that CMS in D male-sterile individuals was associated with a reduction of co-encoded complex I contribution to OXPHOS that might nonetheless be partly compensated by nuclear-encoded complex II. Moreover, the efficiency of the OXPHOS system was similar to N normal hermaphrodite which may explain the female advantage (higher growth) in the absence of energy costs linked to male function. Such changes were not observed in K restored hermaphrodites, suggesting that male fertility restorer genes re-established the OXPHOS metabolism through complex I to a similar level as N normal hermaphrodites. Moreover, the K restored hermaphrodites showed a reduced growth, reflecting the costs of bearing CMS and restorer genes, which may be at least partly explained by the OXPHOS control efficiency and the potential higher relative anaerobic contribution to cellular metabolism that we observed. In the following, we discuss in detail these cellular mechanisms of CMS and their implications for its evolution.

### Effects of CMS on reproductive functions and cellular metabolism in male-steriles

#### Male fertility and female advantage

The conflict opposing the D mitogenome and the nuclear genome results in male sterility in almost all D snails as observed in previous studies (Bererd et al., 2025; David et al., 2022). Less than 2% of D individuals were considered male-fertiles (*i.e.* they fertilised their albino partner), confirming an incomplete penetrance of CMS genes (no restorers are yet found for D mitotype; Bererd et al., 2025; Laugier et al., 2024; Skarlou et al., 2025). The D male-sterile snails had a higher growth, both in terms of mass and length, than N normal hermaphrodite. Bererd et al., (2025) showed that the growth advantage of D male-steriles led to a higher egg production, supporting the expected female advantage in male-sterile individuals required to the maintenance of CMS in natural populations (Dufaÿ et al., 2007). Such a female advantage has been described in plants (Dufay & Billard, 2012) and is usually explained as a consequence of an energy reallocation from the defective male function toward the female fitness (Budar et al., 2003).

#### Complex I impairment and Complex II compensation

Mitonuclear incompatibilities, particularly when mitochondrial and nuclear genomes have not coevolved, may disturb the integrity of OXPHOS with downstream consequences on many cellular functions (Rand et al., 2004; Sunnucks et al., 2017; Wolff et al., 2014). For example, genetically perturbed mitonuclear co-evolved system by introducing a foreign mitogenome from non-human apes (common chimpanzee, pigmy chimpanzee and gorilla) into a human nuclear background resulted in a significant decreased complex I activity and supported mitochondrial respiration, whereas others mitochondrial co-endoded complexes were less affected (Barrientos et al., 1998). Similarly, a foreign mitogenome from rat into a mouse nuclear background resulted in a large decrease of complex I activity, with a lesser impact on co-encoded complex IV activity (McKenzie & Trounce, 2000). Our results are strikingly consistent as the complex I activity *per se* was lower in D mitotype than in N normal hermaphrodite (although not significantly: p = 0.079), whereas complex IV activity remained unaffected. This suggest that CMS in *P. acuta* is associated with an alteration of co-encoded complex I, which likely result from incompatibilities between the extremely divergent D mitogenome and a shared nuclear genome (D snails have to mate with N hermaphrodite snails to produce offsprings; David et al., 2022). Moreover, David et al. (2022) previously showed that the genes coding for NAD proteins (*i.e.* the mitochondrial genes encoding for complex I subunits) exhibited the highest levels of divergence (*e.g.* 54-78% of amino acid dissimilarity for NAD genes of complex I against 19-50% for COX genes of complex IV; see Table S2 for more details). This might explain why the mitonuclear incompatibilities specifically altered the complex I function rather than other co-encoded complexes. Our result is also consistent with studies in CMS plants showing mitochondrial mutations also affecting complex I: tobacco (Gutierres et al., 1997), rice (D. Tang et al., 2021), maize (Marienfeld & Newton, 1994; H. Tang et al., 2017).

Interestingly, although complex I respiration appears impaired in D male-steriles, the combined activity of complexes I and II was similar to N normal hermaphrodites. These results suggest that OXPHOS can offset the dysfunction of complex I by compensating through complex II in D male-sterile individuals (at least, in terms of O_2_ consumption). A similar offset of complex I dysfunction by complex II has been observed in male-steriles in CMS *Nicotina sylvestris,* where alternative NAD(P) H dehydrogenases in association with complex II compensated for the complex I dysfunction, ensuring the plant survival and development (Sabar et al., 2000). As no divergence in the nuclear genes coding for mitochondrial proteins between D male-steriles and N normal hermaphrodites have been reported (David et al., 2022), this suggests that the maintenance of OXPHOS integrity through complex II compensation might not result from an evolutionary rescue by the nuclear genome, but rather might reflect a plastic response of mitochondrial aerobic metabolism. Moreover, we did not detect other compensatory mechanisms: the relative contribution of anaerobic metabolism (LDH/CS ratio) was similar between D male-steriles and N normal hermaphrodites. Consequently, our results suggest that the plasticity of the mitochondrial respiratory chain in response to complex I impairment due to CMS, allowed the maintenance of mitochondrial respiration at the cellular level. This finding may explain the similar whole-body metabolic rate previously observed between D and N snails (Bererd et al., 2025).

#### Consequences of OXPHOS offset on male fertility and female advantage

The hypothesis of a compensation of complex I dysfunction is consistent with a similar OXPHOS control efficiency between D male-steriles and N normal hermaphrodites, suggesting that phosphorylating activity might not be different between mitotypes, which would however need to be confirmed by direct ATP measurement. A maintenance of ATP production in D male-steriles might explain the female advantage conferred by a reallocation of energy from the defective male reproductive function toward growth, thereby enhancing egg production (female fitness; Bererd et al., 2025). However, the dysfunction of complex I of the respiratory chain and the compensation by complex II in D male-sterile individuals could have consequences at the cellular level that may contribute to the sterilization of the male function. Electron entry through complex II into the respiratory chain results in reduced ATP yield and higher reactive oxygen species (ROS) production than electron entry through complex I (Lee et al., 1996; Quinlan et al., 2012). In CMS plant studies, a decrease in ATP production is often proposed to explain the pollen abortion as gametogenesis is highly energy-consuming (Touzet & H. Meyer, 2014). While some examples were consistent with this hypothesis (*e.g.* Li et al., 2012), others did not find such a relationship (*e.g.* Robison et al., 2009; Wen et al., 2003). Similarly, defective ATP production through mitochondrial respiration in animals could lead to disturbed male gametogenesis or spermatozoa performances (Yu et al., 2019) as these cells have limited cytoplasmic space restricting the amount of mitochondria. Previous studies showed that D male-sterile have a very low spermatozoa count, often none at all, indicating rather problems during gametogenesis than lower spermatozoa performances (Bererd et al., 2025; David et al., 2022). Another hypothesis for the mechanisms underlying CMS involves the production of ROS, which may contribute to male sterility (Touzet & H. Meyer, 2014). ROS are molecules that can damage nucleic acids, lipids and proteins, and are involved in cell death (Logan, 2006). In CMS plants, a burst of ROS has been observed in several species (cotton, Jiang et al., 2007; peach, Cai et al., 2021; wheat, Liu et al., 2018) and related to pollen abortion through different mechanisms involving the initiation of cell death process (Touzet & H. Meyer, 2014). In animals, ROS are implicated in reproductive disorders (*e.g.* male infertility in humans; Aitken et al., 2022). Further investigations directly measuring the mitochondrial ATP synthesis and mitochondrial coupling efficiency (*e.g.* ATP/O ratio), as well as the mitochondrial ROS production, antioxidant defences and oxidative damage, would be informative to confirm these hypotheses and are necessary to unravel the role of ATP and ROS production in male sterility.

### Effects of nuclear restoration on reproductive functions and cellular metabolism in restored hermaphrodites

#### Male fertility and costs of CMS and restoration in restored hermaphrodites

The major proportion of K individuals were male fertile, confirming their status of restored hermaphrodites. A previous study provided evidence that nuclear restorer genes counteract the sterilizing effect of CMS genes in the K mitogenome (Laugier et al., 2024). Few K individuals (almost 11%) were considered as male-steriles by our male status test (pairing with a virgin albino partner) but this was likely due to a mating failure independently of the spermatozoa production (see Bererd et al., 2025 for more details). As previously described in Bererd et al. (2025), K restored hermaphrodites showed a lower growth (smaller in mass and length) than N normal hermaphrodites, suggesting that bearing CMS and restorer genes can be costly. Such costs are expected by theoretical models to maintain gynodioecy in populations (*e.g.* restoration costs avoid fixation of nuclear restorers leading to 100% hermaphrodite populations; Dufaÿ et al., 2007). The cost of restoration has been already observed in plants, not in terms of growth impairment, but through detrimental effects on seed (De Haan et al., 1997) or pollen production (Bailey, 2002) or both (Gigord et al., 1999).

#### Normal contribution of Complex I and Complex II to OXPHOS

The mechanisms of male fertility restoration by nuclear genes is complex as it can occur at various molecular levels, from genomic to metabolic levels, and consist in either the inactivation of the sterilizing factor (suppressing the expression of CMS genes) or by offsetting its effects (Bhattacharya et al., 2024 for a recent review). For example, in the CW-CMS rice, the nuclear restorer gene Rf17 encodes for a mitochondrial protein causing metabolic changes and restores the male fertility through poorly known mechanisms (Fujii & Toriyama, 2008). In the *P. acuta* system, our results showed that K restored hermaphrodites had similar complex I and complex II contributions to OXPHOS than in N normal hermaphrodites despite their extremely divergent mitochondrial genomes. Similarly than in D male-sterile snails, the NAD genes (encoding for complex I subunits) in K restored hermaphrodites are the mitochondrial genes with the highest divergence (75-89% of amino acid dissimilarity for NAD genes against 43-70% for COX genes of complex IV; see Table S2 for more details; Laugier et al., 2024). Assuming that mechanism by which CMS genes act to sterilize the male reproductive function in K mitotype are similar than in D male-steriles, our results suggest that nuclear restorer genes in K snails may have restored the complex I function, avoiding the need for complex II compensation to maintain mitochondrial respiration and the associated detrimental consequences for fertility. To confirm that the nuclear restorer genes act on the complex I contribution to OXPHOS, further studies should explore the mitochondrial function of K male-steriles (*i.e.* non-restored), to test for OXPHOS dysfunctions as observed in D male-steriles.

#### Decrease of OXPHOS control efficiency as a cause of CMS and restoration costs

Although complexes I and II contributions to OXPHOS were similar to N normal hermaphrodites, the OXPHOS control efficiency was lower due to a higher proton leak in K restored hermaphrodites. Indeed, a higher mitochondrial proton leak has been shown to limit growth rate in other study systems (*e.g.* Dawson et al., 2022; Salin et al., 2012, 2019). We previously showed that the lower growth of K restored hermaphrodite was not associated with a decrease of egg production or number of spermatozoa (Bererd et al., 2025). Therefore, with the lower amount of ATP available suggested by the lower OXPHOS efficiency, the K restored hermaphrodites are likely unable to maintain the same investment as N normal hermaphrodites in both reproduction and growth, and we can reasonably assume that they thus invest more in reproduction at the expense of growth. The LDH/CS ratio was higher in K mitotype, primarily due to a reduced CS activity rather than an increase in LDH activity. This relative increase in the ratio indicates a higher contribution of the anaerobic pathway to cellular energy metabolism, which is likely to trigger a lower efficient ATP synthesis, unable to meet energy demands in K individuals. To our knowledge, these are the first results showing the cellular mechanisms underlying CMS and restoration costs. However, further investigations will be needed to disentangle the relative effects of CMS and nuclear restorer genes on the energy trade-off between growth and reproduction.

## Conclusion

Our results suggest that CMS was associated with a decreased functionality of complex I in D male-steriles compensated through complex II, allowing an apparently normal mitochondrial respiratory capacity. CMS mechanisms are complex as only male gametes are impaired in D mitotype, while female function remains functional. Similar OXPHOS control efficiency between N hermaphrodites and D male-steriles suggest that ATP production might not differ between these mitotypes, possibly explaining the female advantage from energy reallocation from defective male reproductive function toward growth which indirectly enhanced egg production (Bererd et al., 2025). K restored hermaphrodites showed a similar CI OXPHOS contribution as N normal hermaphrodites, suggesting that restorer genes may restore the complex I function. However, K restored hermaphrodites exhibited a lower OXPHOS control efficiency suggesting a lower capacity to produce ATP and may have a higher contribution of anaerobic metabolism, likely explaining the lower growth associated with the costs of bearing CMS and restorer genes. These results support that mitochondrial aerobic metabolism plays a central role in key parameters underlying the evolutionary dynamics involved in gynodioecy, and positioned the *P. acuta* system as a new model to study the molecular mechanisms of mitonuclear conflicts involved in CMS.

## Acknowledgements

We are grateful to Noéline Garcia for her support with growth measurements and the preparation of individuals for the male sterility test. We also thank Fanny Laugier for conducting the analysis of amino acid divergences among mitotypes. We thank Patrice David for his valuable discussions on cytoplasmic male sterility and for leading the MINIGAN project.

