## Supplementary material for "Cytoplasmic male sterility and mitochondrial metabolism: evidence for low complex I contribution in male-sterile freshwater snail *Physa acuta*"

Table S1. Post-hoc pairwise comparisons between mitotype for mitochondrial respiration rates**.**

| **Respiratory state** | **Comparison** | **Estimate** | **SE** | **df** | **t.ratio** | **p** |
| --- | --- | --- | --- | --- | --- | --- |
| **AOX** | D – K | -0.5563 | 0.205 | 31.6 | -2.715 | **0.0160** |
|  | D – N | -0.5971 | 0.211 | 59.4 | -2.827 | **0.0160** |
|  | K – N | -0.0408 | 0.216 | 41.7 | -0.189 | 0.8511 |
| **CI OXPHOS** | D – K | -0.5255 | 0.205 | 31.6 | -2.565 | **0.0459** |
|  | D – N | -0.4117 | 0.208 | 56.3 | -1.981 | 0.0788 |
|  | K – N | 0.1138 | 0.212 | 38.9 | 0.536 | 0.5953 |
| **CI+II OXPHOS** | D – K | -0.2032 | 0.205 | 31.6 | -0.992 | 0.6956 |
|  | D – N | -0.1195 | 0.208 | 56.3 | -0.575 | 0.6956 |
|  | K – N | 0.0837 | 0.212 | 38.9 | 0.394 | 0.6956 |
| **CI+II LEAK** | D – K | -0.6284 | 0.205 | 31.6 | -3.067 | **0.0132** |
|  | D – N | -0.0465 | 0.208 | 56.3 | -0.224 | 0.8238 |
|  | K – N | 0.5819 | 0.212 | 38.9 | 2.739 | **0.0139** |
| **COX** | D – K | 0.0648 | 0.205 | 31.6 | 0.316 | 0.9185 |
|  | D – N | -0.0214 | 0.208 | 56.3 | -0.103 | 0.9185 |
|  | K – N | -0.0862 | 0.212 | 38.9 | -0.406 | 0.9185 |
| **ETS** | D – K | -0.1034 | 0.205 | 31.6 | -0.505 | 0.9177 |
|  | D – N | -0.1255 | 0.208 | 56.3 | -0.604 | 0.9177 |
|  | K – N | -0.0221 | 0.212 | 38.9 | -0.104 | 0.9177 |

Table S2. Divergence rate of protein sequence between mitotypes D male-steriles and N normal hermaphrodites, and K restored hermaphrodites and N**.**

| Gene | Complex | D_vs_N | K_vs_N |
| --- | --- | --- | --- |
| ATP6 | V | 0.698 ± 0.008 | 0.714 ± 0.007 |
| COX1 | IV | 0.427 ± 0.002 | 0.377 ± 0.001 |
| COX2 | IV | 0.677 ± 0.006 | 0.671 ± 0.005 |
| COX3 | IV | 0.697 ± 0.006 | 0.668 ± 0.003 |
| CYTB | III | 0.64 ± 0.007 | 0.624 ± 0.004 |
| NAD1 | I | 0.75 ± 0.007 | 0.733 ± 0.008 |
| NAD2 | I | 0.835 ± 0.008 | 0.834 ± 0.004 |
| NAD3 | I | 0.778 ± 0.015 | 0.756 ± 0.011 |
| NAD4 | I | 0.768 ± 0.002 | 0.768 ± 0.002 |
| NAD5 | I | 0.783 ± 0.003 | 0.699 ± 0.006 |
| NAD6 | I | 0.889 ± 0.011 | 0.82 ± 0.011 |

Figure S1. **Mitochondrial respiration rates** for each state. Colored dots represent individual values. Black dots and error bars correspond to means ± standard error of raw data.

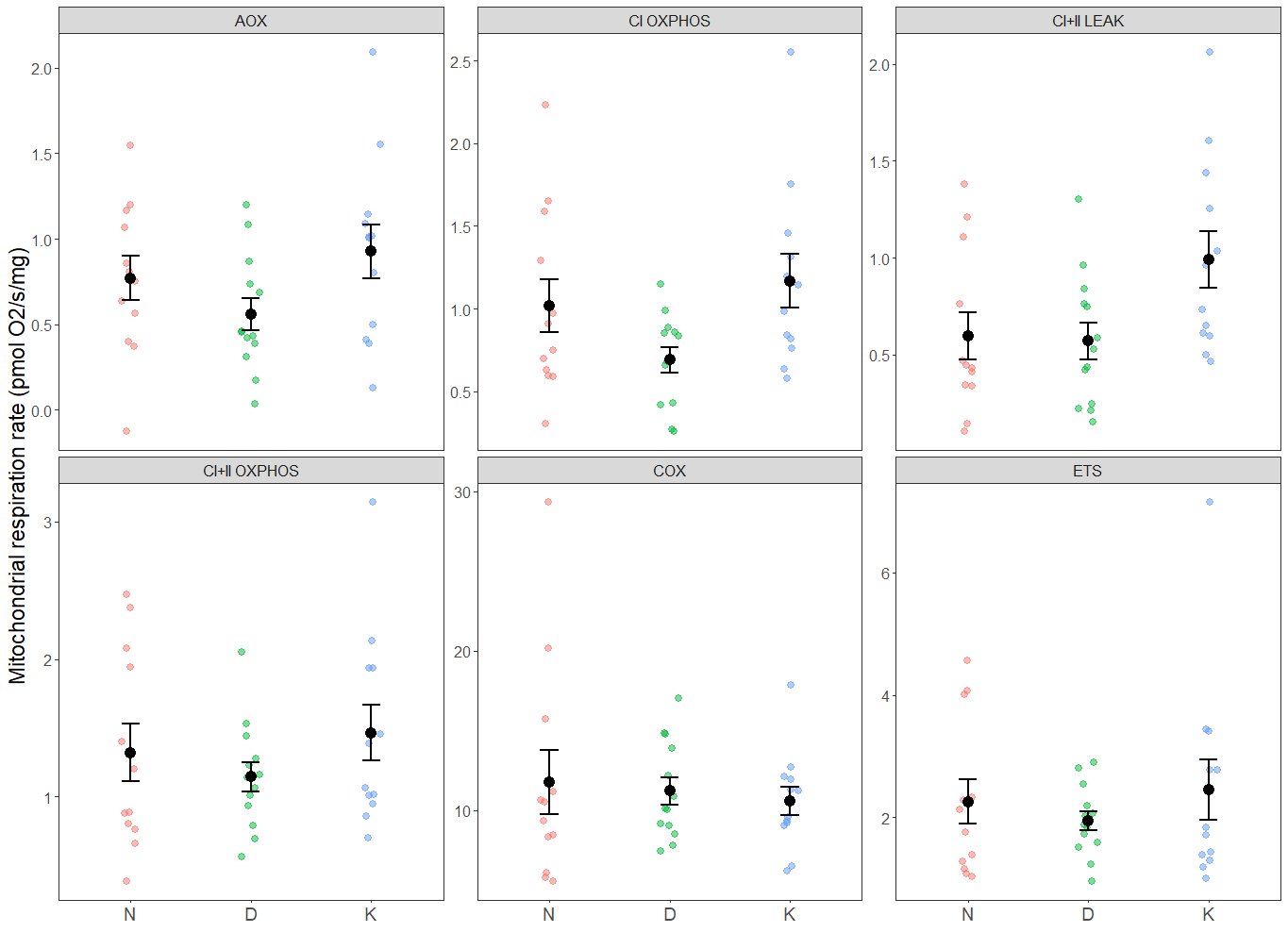

Figure S2. **Mitochondrial control ratio**. Colored dots represent individual values. Black dots and error bars correspond to means ± standard error of raw data.

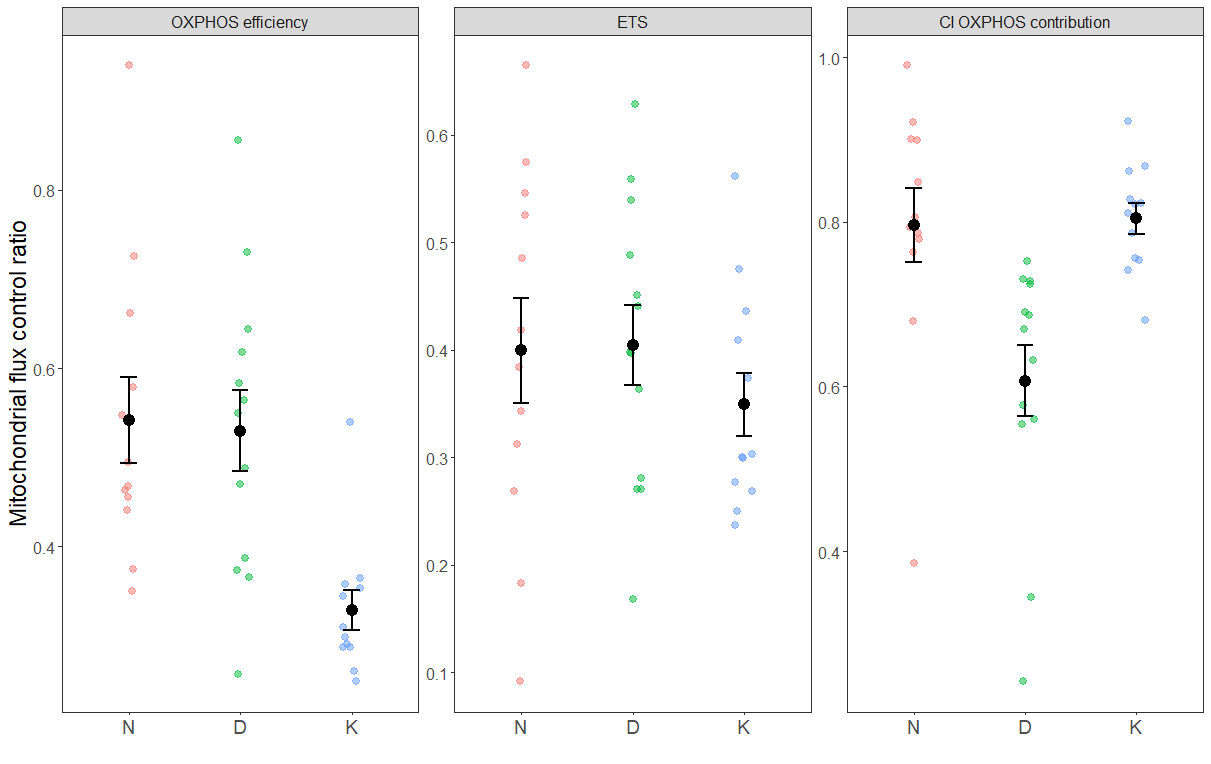

Figure S3. **Enzymatic activities**. Colored dots represent individual values. Black dots and error bars correspond to means ± standard error of raw data.

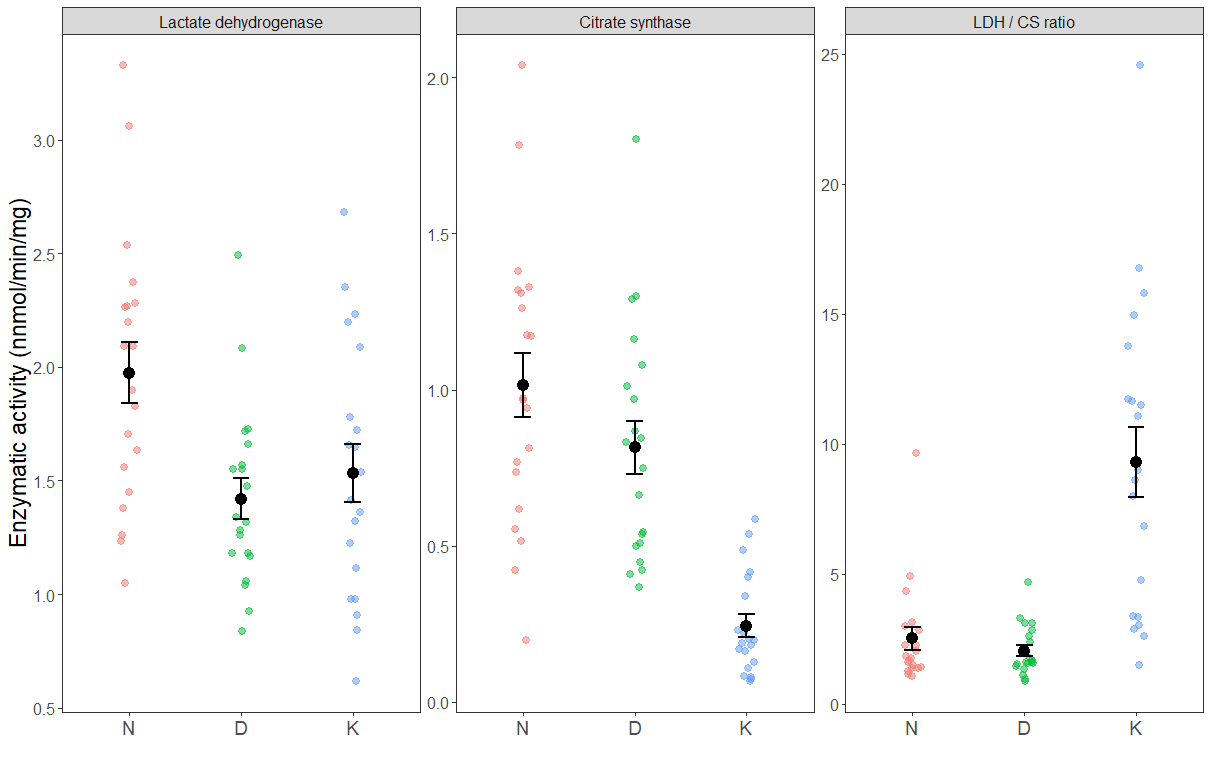
